# Acute treatment with TrkB agonist LM22A-4 confers neuroprotection and preserves myelin integrity in a mouse model of pediatric traumatic brain injury

**DOI:** 10.1101/2020.10.01.321570

**Authors:** Jessica L. Fletcher, Larissa K. Dill, Rhiannon J. Wood, Sharon Wang, Kate Robertson, Simon S. Murray, Akram Zamani, Bridgette D. Semple

**Affiliations:** Department of Anatomy and Neuroscience, University of Melbourne, Parkville, VIC Australia; Department of Neuroscience, Central Clinical School, Monash University, Melbourne, VIC Australia; Department of Medicine (Royal Melbourne Hospital), The University of Melbourne, Parkville, VIC, Australia

**Keywords:** Tropomyosin-related kinase receptors, LM22A-4, brain-derived neurotrophic factor, BDNF, myelination, inflammation, external capsule, corpus callosum, spectral confocal reflectance microscopy

## Abstract

Young children have a high risk of sustaining a traumatic brain injury (TBI), which can have debilitating life-long consequences. Importantly, the young brain shows particular vulnerability to injury, likely attributed to ongoing maturation of the myelinating nervous system at the time of insult. Here, we examined the effect of acute treatment with partial tropomyosin receptor kinase B (TrkB) agonist, LM22A-4, on the pathological and neurobehavioral outcomes after pediatric TBI, with the hypothesis that targeting TrkB would minimize tissue damage and support functional recovery. We focused on myelinated tracts— the corpus callosum and external capsules—based on recent evidence that TrkB activation potentiates oligodendrocyte remyelination. Male mice at postnatal day 21 received an experimental TBI or sham surgery. Acutely post-injury, extensive cell death, a robust glial response and disruption of compact myelin were evident in the injured brain. TBI or sham mice then received intranasal saline vehicle or LM22A-4 for 14 days. Behavior testing was performed from 4 weeks post-injury, and brains were collected at 5 weeks for histology. TBI mice showed hyperactivity, reduced anxiety-like behavior, and social memory impairments. LM22A-4 ameliorated the abnormal anxiolytic phenotype but had no effect on social memory deficits. Use of spectral confocal reflectance microscopy detected persistent myelin fragmentation in the external capsule of TBI mice at 5 weeks post-injury, which was accompanied by regionally distinct deficits in oligodendrocyte progenitor cells and postmitotic oligodendrocytes, as well as chronic reactive gliosis and atrophy of the corpus callosum and injured external capsule. LM22A-4 treatment ameliorated myelin deficits in the perilesional external capsule, as well as tissue volume loss and the extent of reactive gliosis. However, there was no effect of this TrkB agonist on oligodendroglial populations detected at 5 weeks post-injury. Collectively, our results demonstrate that targeting TrkB immediately after TBI during early life confers neuroprotection and preserves myelin integrity, and this was associated with some improved neurobehavioral outcomes as the pediatric injured brain matures.

## INTRODUCTION

Traumatic brain injury (TBI) is a leading cause of death and disability worldwide (Maas et al., 2008; Thurman, 2016), and in children under 5 years of age, it can lead to life-long neurological dysfunction that may not manifest until adolescence and adulthood (Di Battista et al., 2012; Ryan et al., 2016). The emergence of these long-term psychosocial and behavioral impairments is attributed to the coincident timing of pediatric TBI with a highly active period of postnatal brain development that includes synaptogenesis, synaptic pruning, and rapid developmental acquisition of myelin (Semple et al., 2013). In particular, the extent and persistence of damage to myelinated white matter tracts, including the corpus callosum—the connection between left and right hemispheres—is thought to be a major causal contributor to neurobehavioral dysfunction in pediatric TBI, based on natural history studies using magnetic resonance (MRI) / diffusion tensor imaging (DTI) to track injury-induced neuropathology (Ewing-Cobbs et al., 2008; Oni et al., 2010; Wilde et al., 2012; Wu et al., 2010). However, the cellular and molecular pathology that these MRI / DTI measures represent to axons, their myelin sheaths and the oligodendroglia that produce myelin, and how the developmental trajectory of these cellular structures are affected by early life TBI, are poorly understood.

Preserving the structural integrity of myelinated tracts, and targeting the process of myelination immediately after a pediatric TBI, could be a novel therapeutic avenue to prevent the manifestation of neurobehavioral and social problems in adulthood that severely impacts quality of life (Shi et al., 2015). Indeed, recent evidence that specifically targeting the activation of oligodendroglial-expressed TrkB-receptors promotes myelin repair in the context of adult cuprizone-induced demyelination (Fletcher et al., 2018a; Nguyen et al., 2019) suggests that such remyelinating therapies may also yield benefit in the context of TBI. One such potential therapy is the promising small molecule partial TrkB agonist LM22A-4. This compound was originally identified for its potential to emulate the Loop IIb subregion of brain-derived neurotrophic factor (BDNF) (Massa et al., 2010), which is required to trigger TrkB autophosphorylation and engage downstream intracellular signaling cascades PI3K/Akt, PLCγ and MAPK/Erk (Chao, 2003). The latter is essential for the pro-myelinating activity of BDNF-TrkB signaling (Du et al., 2006; Van’t Veer et al., 2009; Xiao et al., 2012). LM22A-4 also has neuroprotective properties, and has been reported to promote functional motor recovery in rodent models of adult TBI, Huntington disease and stroke (Han et al., 2012; Simmons et al., 2013), as well as neuronal survival in BDNF-dependent hippocampal cultures (Massa et al., 2010).

In the present study, we tested the hypothesis that LM22A-4 treatment would preserve myelin integrity and promote neuroprotection in a mouse model of moderately-severe experimental TBI. This model represents toddler aged children, and corresponds with a highly active developmental period in central nervous system (CNS) myelin acquisition (Eakin et al., 2014; Emery et al., 2009; Hill and Grutzendler, 2019; Hill et al., 2018). Acute treatment was performed to maximize the likelihood of mitigating future cognitive, social and behavioral consequences of pediatric TBI (McCarthy et al., 2006), while intranasal delivery was chosen to rapidly increase the accumulation of LM22A-4 directly into the cerebrospinal fluid and CNS while bypassing the bloodstream (Born et al., 2002). Collectively, our findings demonstrate that acute intervention with a partial TrkB agonist can preserve myelin integrity and improve some of the aberrant adolescent neurobehavioral outcomes after pediatric TBI.

## MATERIALS AND METHODS

### Animals and experimental groups

Male C57BL/6J mice were obtained from the on-site breeding colony at the Florey Institute of Neuroscience and Mental Health animal facility in Melbourne, generated from breeding pairs purchased from the Animal Resource Centre (Perth, Australia). Only males were used for the current study, as even at a young age, males are more likely to sustain a TBI (Thurman, 2016). Mice were housed in a specific-pathogen-free facility, under a 12 hour light/dark cycle with *ad libitum* access to food and water. All procedures were performed with approval from the institutional Animal Ethics Committee (#16-101-UM) and in accordance with the National Health and Medical Research Council (NHMRC) Australian Code of Practice for the Care and Use of Animals for Scientific Purposes.

To examine acute pathology after pediatric TBI, mice were randomized to receive either TBI or sham surgery, then euthanized at 3 days (n=3/group). To confirm TrkB activation by the agonist LM22A-4, mice were randomized to receive either TBI or sham surgery, followed by randomization to treatment with the TrkB agonist LM22A-4 or 0.9% saline vehicle control as described below. Animals were euthanized at 2 weeks post-injury for Western blot analyses (n=3/group). To examine the effect of TrkB agonism on outcomes after pediatric TBI, mice were randomized to receive either TBI or sham surgery, followed by treatment with the TrkB agonist LM22A-4 or vehicle control. These animals were euthanized at 5 weeks post-injury following a period of behavior testing (n=6/group; see <u>Supplementary Table 1</u>). Group allocation was coded to ensure that investigators were blinded during data collection and analysis, as per best practice guidelines (Landis et al., 2012).

### Experimental TBI

At postnatal day 20-22, mice were randomized to either the controlled cortical impact model of experimental TBI or sham surgery, as previously described (Semple et al., 2017; Tong et al., 2002). Briefly, mice were anesthetized with isoflurane and positioned in a stereotaxic frame, then administered 0.05 mg/kg s.c. buprenorphine in isotonic saline. The skull was exposed by midline incision, and a micro-drill was used to generate a 4 mm diameter circular craniotomy over the left parietal lobe, midway between Bregma and Lambda. The impact to the intact dura was induced by a custom-built computer-controlled cortical impactor device with the following settings for a moderately-severe TBI: 3.0 mm tip; velocity 4.5 m/s; depth 1.7 mm; duration 150 ms. Sham mice underwent identical surgical procedures with the exception of the cortical impact. To complete the surgery, the scalp was closed with sutures, animals were removed from anesthesia and allowed to recover in heated chambers for ~ 1 h under close observation, before being returned to group housing. Body weights were monitored daily for 1 week then weekly thereafter until euthanasia.

### LM22A-4 administration

The TrkB agonist LM22A-4 (Massa et al., 2010) was purchased from Abcam (ab22020) and reconstituted to a 100μg/μL stock, then diluted to 10μg/μL in 0.9% isotonic saline for use. LM22A-4 or vehicle control (saline only) was administered to awake mice via the intranasal route at 5 mg/kg, commencing immediately post-injury/sham surgery, then once daily for 2 weeks.

### Behavior testing

At 4 weeks post-injury, mice (n=6/group) were individually housed for a week of behavior testing, conducted within the Florey Institute of Neuroscience and Mental Health animal facility in allocated mouse behavior rooms. Assessments were performed in the following order: Open Field, Elevated Plus Maze, rotarod, three-chamber social approach test. All testing was conducted by investigators blinded to injury and treatment groups.

General activity was evaluated in the open field paradigm (Semple et al., 2017). Experimental mice were allowed a 10 min period of free exploration of the testing arena (33 x 33 x 27 cm). Movements were tracked by an overhead camera and TopScan software (Clever Sys Inc, Reston, VA) as a measure of general activity.

The Elevated Plus Maze was used to evaluate anxiety-like behavior, based on the tendency of rodents to avoid open spaces (Semple et al., 2017). During a 10 min period, mice were permitted free exploration of the apparatus, which consists of 2 open arms and 2 enclosed arms (30 cm x 6 cm) (San Diego Instruments, San Diego, USA). Distance moved and time spent in the open arms, as a percentage of total time, was quantified by live video streaming from an overhead camera using TopScan tracking software.

The three-chamber paradigm is the gold standard test to evaluate same-sex social affiliation and social recognition in mice (Yang et al., 2011). The task was performed as described previously (Semple et al., 2012), with some modifications. The apparatus was 42 x 39 cm, with a central chamber of 9 cm diameter and two outer chambers of 16.5 cm diameter. Same-sex, age-matched, naive stimulus mice were restricted to rectangular metal cages (11 x 16 cm) fitted at one end of the outer chambers, such that experimental mice could only approach and initiate a social investigation from one side. Testing consisted of three consecutive 10 minute stages: (1) habituation in the chamber, containing two empty cages; (2) a choice between an empty cage and a cage containing a stimulus mouse; and (3) a choice between a second ‘novel’ stimulus mouse and the first, now ‘familiar’ mouse. Stages 2 and 3 are considered measures of the subject’s preference for sociability and social recognition/social memory, respectively.

Stimulus mice and their relative positioning (left versus right chambers) were randomized between subjects. Data were expressed as time spent in each of the outer chambers as a percentage of total time.

### Tissue collection

All mice were euthanized with sodium pentobarbital, 80 mg/kg i.p. For histological analyses, mice were transcardially perfused with 0.1 M sterile isotonic phosphate buffered saline (PBS) followed by 4% paraformaldehyde (PFA). Brains were excavated from the skull, post-fixed overnight in 4% PFA, then cryopreserved in 30% sucrose prior to embedding in optimal cutting temperature medium. Frozen samples were cryosectioned in the coronal orientation at 12 μm thickness from Bregma −0.9 mm to approximately −3.3 mm, and collected onto Superfrost Plus slides (Grale Scientific), for storage at −80 °C until use.

For Western blotting, mice were transcardially perfused with 0.1M isotonic PBS before decapitation to remove the brain. The cerebral hemispheres and bilateral olfactory bulbs were dissected, snap frozen and homogenized for Western blotting to detect total and phosphorylated TrkB and ERK1/2 (see <u>Supplementary Methods</u>).

### Immunofluorescence staining

Immunofluorescence staining was performed as previously described (Fletcher et al., 2018b; Webster et al., 2019). Briefly, slides were air-dried and rehydrated in PBS, before overnight incubation at room temperature or 4 °C with primary antibodies (Table 1). Slides were then washed, incubated in blocking solution containing species-specific normal serum and 0.1% Triton X-100, and incubated with the appropriate fluorophore-conjugated secondary antibody for 1-2 h at room temperature in the dark. Slides were washed, and counterstained with the nuclear marker Hoescht 33442, before mounting with aqueous DAKO fluorescence mounting media. All immunohistochemistry was performed in batches (mixed experimental groups) containing a negative control slide (primary antibodies omitted).

**Table 1:** Primary antibodies used for immunofluorescence staining of brain tissue sections.

| Antibody | Used to identify | Vendor, Code | Concentration |
| --- | --- | --- | --- |
| Rat anti-myelin basic protein (MBP) | Myelin | Millipore, MAB386 | 1:200 |
| Rabbit polyclonal anti-beta amyloid precursor protein (β-APP) | Axonal disruption | Life Technologies, 51-2700 | 1:500 |
| Goat polyclonal anti-ionized calcium-binding adaptor protein (Iba1) | Microglia and macrophages | Abcam, Ab5076 | 1:750 |
| Rabbit polyclonal anti-glial fibrillary acidic protein (GFAP) | Astrocytes | DAKO, Z0334 | 1:1000 |
| Mouse monoclonal anti-CCI | Post-mitotic oligodendrocytes | CalBioChem, OP80 | 1:200 |
| Rabbit polyclonal anti-Olig2 | Oligodendrocyte lineage | Millipore, AB9610 | 1:200 |
| Polyclonal goat anti-platelet-derived growth factor receptor alpha (PDGFRα) | Oligodendrocyte precursor cells (OPCs) | R&D Systems, AF1062 | 1:200 |

GFAP and Iba1 were double-labeled, to detect reactive astrocytes and microglia, respectively. CC1 (APC), Olig2 and PDGFRα were stained using a triple labeling protocol, to detect postmitotic mature oligodendrocytes (CC1^+^), oligodendrocyte precursors (PDGFRα^+^) and the entire oligodendrocyte lineage (Olig2^+^), with all antibodies incubated simultaneously.

Secondary antibodies were donkey anti-rat IgG Alexa Fluor (AF) 488 (for MBP); donkey antirabbit IgG AF 594 (for β-APP or Olig2); donkey anti-goat IgG AF 488 (for Iba1 or PDGFRα); donkey anti-mouse IgG AF 488 (for GFAP); or donkey anti-mouse IgG AF647 (for CC1) (1:200 or 1:250; Molecular Probes). All slides were counterstained with Hoechst dye to identify cell nuclei. For MBP, slides were additionally permeabilized by fixation with ice cold 100% methanol prior to blocking and antibody incubations, and the Hoechst counterstain was omitted. For GFAP/Iba-1 and β-APP, slides were also incubated in 0.3% Sudan Black prior to cover-slipping to reduce tissue auto-fluorescence.

Lastly, terminal deoxynucleotidyl transferase dUTP nick end labeling (TUNEL) was performed using the *In Situ* Cell Death Detection Kit, Fluorescein (Roche, #11684795910), as per the manufacturer’s protocol.

### Fluorescence imaging and analysis

A Nikon Ti-E fluorescence microscope with a sCMOS Ando Zyla camera (Monash Micro Imaging facility, Alfred Research Alliance) was used to visualize most of the fluorescence staining (MBP, βAPP, TUNEL, GFAP and Iba1). Photomicrographs of the key regions of interest (ROI)—medial corpus callosum, ipsilateral and contralateral external capsule—were acquired at 10 or 20x magnification using consistent exposure times via NIS Elements Advanced Research software (Nikon). Regions were defined based upon Hoechst labeling or phase-contrast images of the same field of view. For each analysis, a minimum of 4 equidistant sections per animal through the injured brain were used, and data were expressed as an average percentage area of positive staining in a single ROI.

Olig2/PDGFRα/CC1 staining of oligodendroglia was visualized and imaged on a Nikon A1r confocal microscope with 405, 488, 561 and 640 nm laser lines using a Plan Fluro 20x MI 0.75 NA WD 0.66mm glycerin immersion objective and NIS Elements Advanced Research software (Nikon), in the Monash Micro Imaging facility. Tile scans of the medial corpus callosum, ipsilateral and contralateral external capsule were captured in 12 μm thick z-stacks using consistent imaging settings. All cell counts were performed manually by an investigator blinded to experimental group, using FIJI/ImageJ (http://imagej.nih.gov/ij/; National Institutes of Health). Data are expressed as density (cells/mm^3^).

### Spectral confocal reflectance (SCoRe) imaging and analysis

Spectral confocal reflectance (SCoRe) microscopy was used to detect compact myelin (Gonsalvez et al., 2019; Schain et al., 2014). This label-free imaging technique was performed on fixed coronal sections of the medial corpus callosum, ipsilateral and contralateral external capsule, using a Zeiss LSM880 confocal microscope with a Zeiss 40X/1.0 NA water immersion objective, at the Biological Optical Microscopy Platform hosted by the University of Melbourne. Lasers with wavelength 488, 561 and 633 nm were passed through an Acousto-Optical Tunable Filter and a 39 dichroic mirror MBS T80/R20 with the reflected light captured by three photodetectors, set to collect light ± 5 nm around each laser wavelength using prism and mirror sliders. For analysis, the three channels were merged to form a composite greyscale image. The settings for image acquisition remained constant for all samples. To measure compact myelin, the percentage area positive for SCoRe signal was measured using the threshold function in FIJI as previously described (Gonsalvez et al., 2019; Govier-Cole et al., 2019), in restricted ROIs of 100 × 100 μm^2^ in the corpus callosum, contralateral and ipsilateral external capsules, on three sections per brain spaced approximately 144 μm apart from approximately −1.7 mm to −2.6 mm Bregma.

### Volumetric analysis of cresyl violet staining

Five equidistant sections per brain (360 μm apart), adjacent to those used for immunofluorescence staining, were allocated for cresyl violet staining and subsequent volumetric estimation of the corpus callosum, ipsilateral and contralateral external capsule, cortex and hippocampus. Slides were stained with cresyl violet acetate for 15 min followed by differentiation in descending ethanol concentrations as described previously (Semple et al., 2017). One set of slides was additionally stained with Luxol Fast Blue Images to differentiate myelinated tracts, while a second adjacent set of slides were stained with cresyl violet only. Images were captured using the Leica Aperio AT Turbo Brightfield slide scanner within the Monash Histology Platform, then exported to FIJI for analysis using the unbiased Cavalieri method with grid point counting, whereby volume equates to the number of points counted multiplied by the area represented per point and the distance between sections, taking into account sampling frequency and section thickness (Webster et al., 2019). Measurements were restricted to the dorsal hemispheres defined by an inferior horizontal boundary line perpendicular to the most ventral point of the third ventricle at the midline. Different grid sizes were used for different ROIs depending on their size and shape to ensure sufficient points were counted for accurate estimates, as confirmed by co-efficiencies of error averaging <0.05.

### Statistical analyses

Tissue volume, GFAP and Iba1 immunostaining and behavioural data were analyzed using GraphPad Prism v.7. Multivariate analyses of variance (ANOVAs) were performed, with independent variables of treatment and injury, and a within-subject (repeated) dependent variable of time as appropriate. *A priori*, multiple comparison post-hoc tests were used following significant main effect interactions and are annotated graphically where significant. SCoRe and oligodendroglial cell count data were analyzed by restricted maximum likelihood (REML) mixed models (GAMLj package v.2.0.6) (Gallucci, 2020), with estimated distance from impact site, injury and treatment as fixed effects and animal identity as the random cluster variable in Jamovi (v1.1.9). Statistical significance was set as p<0.05, and data are presented as group means with standard error of the mean (SEM).

## RESULTS

### Pediatric TBI results in acute pathology in the injured cortex and underlying myelinated tracts

We first characterized the acute effects of pediatric TBI in a mouse model, focusing on myelinated tracts, by examining brain sections collected at 3 days post-injury. TBI resulted in an obvious lesion core, evident by loss of cresyl violet cellular staining within the posterior parietal lobe, including the somatosensory cortex (Figure 1a). The lesion epicenter was filled with Iba1^+^ monocyte-derived cells (Figure 1b); likely reflecting a combination of peripherally-derived infiltrating macrophages and activated microglia. GFAP immunoreactivity, indicative of reactive astrogliosis, surrounded the lesion core as well as the underlying myelinated tract of the external capsule and the ipsilateral hippocampus. TUNEL-positive, dead/dying cells were abundant in the lesion core and external capsule (Figure 1c), while the accumulation of βAPP confirmed axonal transport disruption in the external capsule (Figure 1d). The external capsule and corpus callosum were observed to be in an active myelinating phase at this age, as evident from SCoRe microscopy which revealed sparse levels of compact myelin in these regions. However, compact myelin debris was evident within the ipsilateral external capsule and perilesional cortex (Figure 1e). Finally, an increase in phosphorylated TrkB (pTrkB) levels was observed in cells proximal to the lesion core, with the majority appearing to have a neuronal cell morphology (Figure 1f).

**Figure 1:**
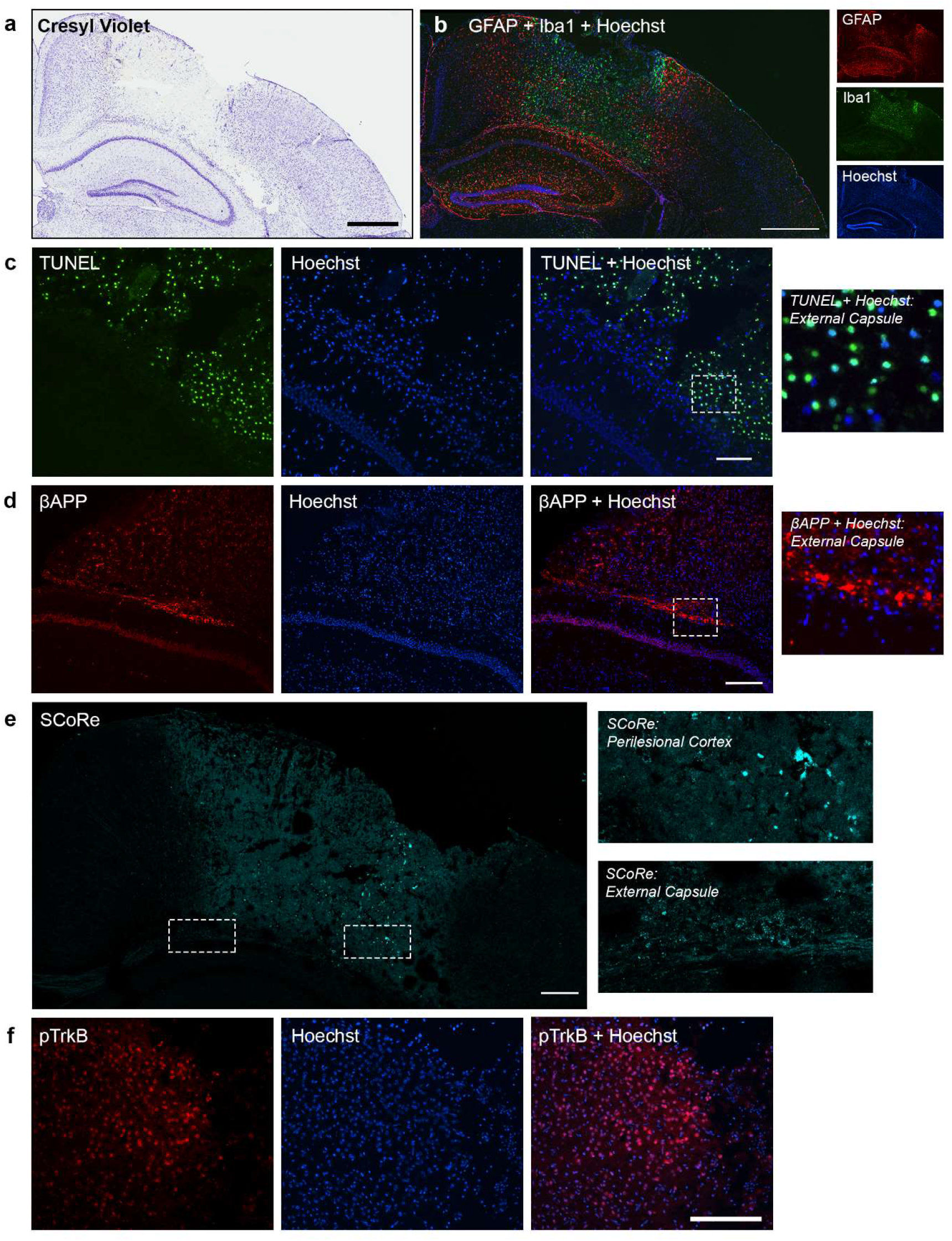
Pediatric TBI in the mouse results in acute pathology in the ipsilateral cortex and underlying myelinated tracts. Representative micrographs illustrating the extent of gross pathology at 3 days post-injury, showing (a) loss of cresyl violet staining, and (b) a glial response around the lesion site (scale bar = 600 μm). GFAP and Iba1 immunostaining indicate reactive astrocytes and microglia/macrophages, respectively. Abundant cell death was detected by (c) TUNEL in the cortex and external capsule (scale bar = 100 μm). Damage to myelinated tracts included accumulation of (d) βAPP in the external capsule, and (e) disruption of compact myelin including myelin debris as detected by SCoRE imaging (scale bar = 200 μm). (f) Intracellular phosphorylation of TrkB was also evident acutely after pediatric TBI (scale bar = 200 μm).

### Intranasal LM22A-4 administration does not influence post-injury weight gain

Activating TrkB signaling has been shown to enhance cellular measures of myelin repair (Fletcher et al., 2018b; Nguyen et al., 2019) and improve motor function outcomes in an adult TBI mouse model (Massa et al., 2010). To determine if this strategy would have beneficial effects in the context of pediatric TBI, the partial TrkB agonist LM22A-4 (0.5 mg/kg) was intranasally administered daily for 14 days. Western blotting was performed on olfactory bulb samples, collected 1 h following the final treatment administration, to assess whether LM22A-4 induced TrkB signaling via the intranasal route (<u>Supplementary Figure 1</u>). This revealed a trend towards increased TrkB phosphorylation in the olfactory bulb of mice treated with LM22A-4 compared to vehicle (2-way ANOVA effect of treatment, F_1,8_=4.57, p=0.06), regardless of injury (F_1,8_=0.23, p=0.64); suggestive of successful induction of TrkB activity by intranasal delivery.

Body weights were recorded as a measure of general animal health over time, and to address the potential that LM22A-4 may affect appetite and influence body weights as suggested previously (Rios, 2013; Waterhouse and Xu, 2013). Over time post-surgery, all groups demonstrated an increasing body weight as anticipated with increasing age (2-way ANOVA effect of time, F_218,360_=574.3, p<0.0001), but no detectable influence of TBI surgery or LM22A-4 treatment (F3, 20=0.51, p=0.68; <u>Supplementary Figure 2</u>). This lack of effect of LM22A-4 on body weights is consistent with previous studies utilizing this partial TrkB agonist over long durations in experimental models (Simmons et al., 2013).

### TBI-induced dysmyelination in the ipsilateral external capsule is limited by acute intranasal LM22A-4 administration

As noted above, compact myelin debris was evident at 3 days post-TBI in the injured cortex and external capsule (Figure 1e). To examine the consequences of this acute pathology more chronically, the medial corpus callosum and ipsilateral external capsule were examined at 5 weeks post-injury. In contrast to the 3 day injured brain at age p23-25, there was minimal βAPP^+^ immunoreactivity in the external capsule of either vehicle or LM22A-4 treated TBI-mice at 5 weeks post-injury (age p58-60) (<u>Supplementary Figure 3</u>). This is consistent with the minimal levels of βAPP^+^ axonal dystrophy within 48 h for younger (p7) animals (Dikranian et al., 2008), or at 7 days after experimental TBI in adult rodents (Weber et al., 2019).

We next examined disruption to myelinated axons, by immunostaining for the major myelin constituent myelin basic protein (MBP), and using spectral confocal reflectance (SCoRe) microscopy to detect compact myelin (Gonsalvez et al., 2019; Schain et al., 2014). At 5 weeks post-injury, MBP immunofluorescence (<u>Supplementary Figure 4</u>) and SCoRe imaging (Figure 2) revealed fragmentation and disruption to fiber tract directionality and continuity after TBI in vehicle-treated animals, that extended from the ipsilateral external capsule immediately ventral to the impact site to the medial corpus callosum, laterally adjacent. However, rarely was there complete absence of MBP or SCoRe signal indicative of profound myelin loss. Interslide variability in the intensity of MBP staining, likely due to increased antigenicity of MBP after injury (Gonsalvez et al., 2019), rendered MBP immunofluorescence inappropriate to quantify changes in myelin after TBI. Instead, the area positive for SCoRe signal was measured using series of restricted ROIs from approximately 0.56 mm rostral to the injury site to approximately 0.14 mm caudal from the injury site in the corpus callosum, and 0.28 mm caudal in the ipsilateral external capsule. A rostro-caudal gradient was observed in the percentage area of compact myelin in the corpus callosum, with a reduction at the site of impact in sham mice only, irrespective of LM22A-4 treatment (REML Mixed Model, location × injury interaction, F_1,5_=3.26, p=0.007; Figure 2b). In mice that received a TBI, this rostro-caudal variation was not detected (Bonferroni’s post-hoc p=0.99).

**Figure 2:**
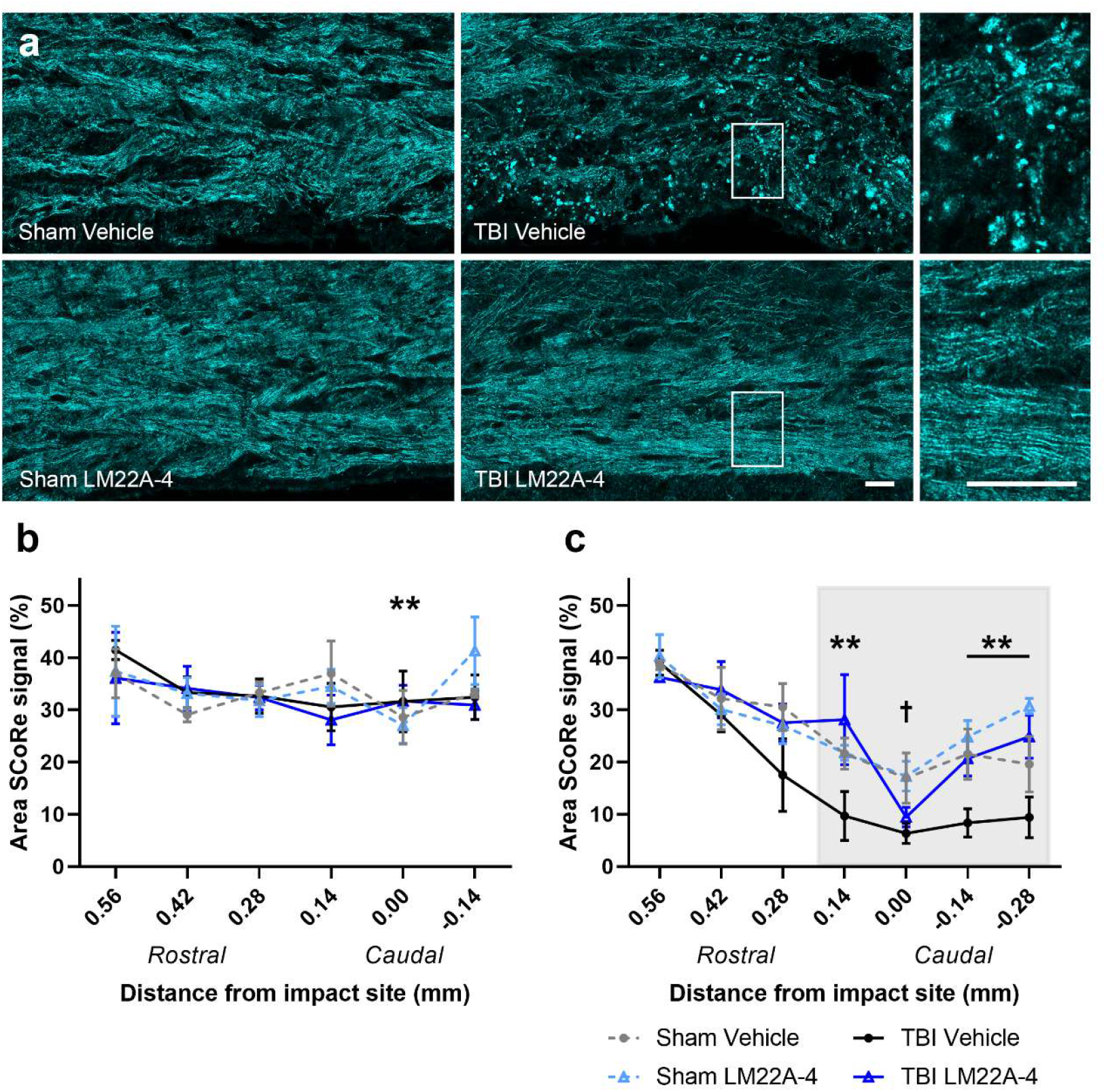
Accumulation of TBI-induced myelin fragmentation and dysmyelination is prevented by acute intranasal treatment with LM22A-4. Representative SCoRe micrographs of the ipsilateral external capsule (a), within the impact zone, showing intact compact myelin in the sham-vehicle, sham-LM22A-4 and TBI-LM22A-4 treated mice, but disrupted and fragmented compact myelin debris in TBI-vehicle mice at 5 weeks post-injury (scale bars = 20 μm). The area of intact compact myelin in the medial corpus callosum (b) was reduced at the impact site in sham mice compared to more rostral regions (**post-hoc p<0.01), indicating normal rostro-caudal variation. A loss of compact myelin in the ipsilateral external capsule (c) of TBI-vehicle mice was observed in all perilesional areas (gray; **post-hoc p=0.002), which was prevented by intranasal LM22A-4 treatment except at the direct impact site (sham vs. TBI: †post-hoc p=0.013). For (b-c), mean ± SEM plotted, n=6/group, REML, Bonferroni adjusted p-values used for post-hoc comparisons.

In the external capsules, a rostro-caudal gradient in compact myelin was also observed (Figure 2c), with reduced SCoRe signal after 0.28 mm rostral from the impact site (ipsilateral: REML Mixed Model, location × treatment × injury interaction, F_1,6_=3.02, p=0.007). In the ipsilateral external capsule, TBI further reduced the percentage area of SCoRe signal across the lesion core and adjacent perilesional areas (Bonferroni’s post-hoc p=0.002 and p<0.001, respectively; Figure 2c). LM22A-4 treatment prevented the perilesional loss of compact myelin. In TBI mice treated with LM22A-4, the percentage area of SCoRe signal in perilesional areas were similar to compact myelin levels detected at these sites in sham-vehicle (Bonferroni’s post-hoc p=0.145) and sham-LM22A-4 control mice (Bonferroni’s post-hoc p=0.99). However, at the level of the impact site, compact myelin was not preserved in LM22A-4 treated mice compare to mice in sham control groups (Bonferroni’s post-hoc p=0.013; Figure 2c), likely due to profound tissue loss remaining in LM22A-4 treated TBI-mice. Collectively, these results indicate that the acute intervention with the partial TrkB agonist, LM22A-4, can preserve (or repair) perilesional myelinated tracts towards healthy levels.

### Regionally distinct losses of oligodendrocyte progenitor cell (OPCs) and post-mitotic oligodendrocytes persist for up to 5 weeks post-TBI

We next examined the oligodendroglial populations in the corpus callosum and ipsilateral external capsule at 5 weeks post-TBI, using triple-immunostaining to co-label the oligodendroglial lineage marker Olig2, with the post-mitotic oligodendrocyte marker APC/CC1 (Figure 3a) or the oligodendrocyte progenitor cell (OPC) marker PDGFRα (Figure 3e). Given that the disruption to compact myelin was highly localized to the impact site and perilesional areas, our assessments of oligodendroglia were focused on these regions.

**Figure 3:**
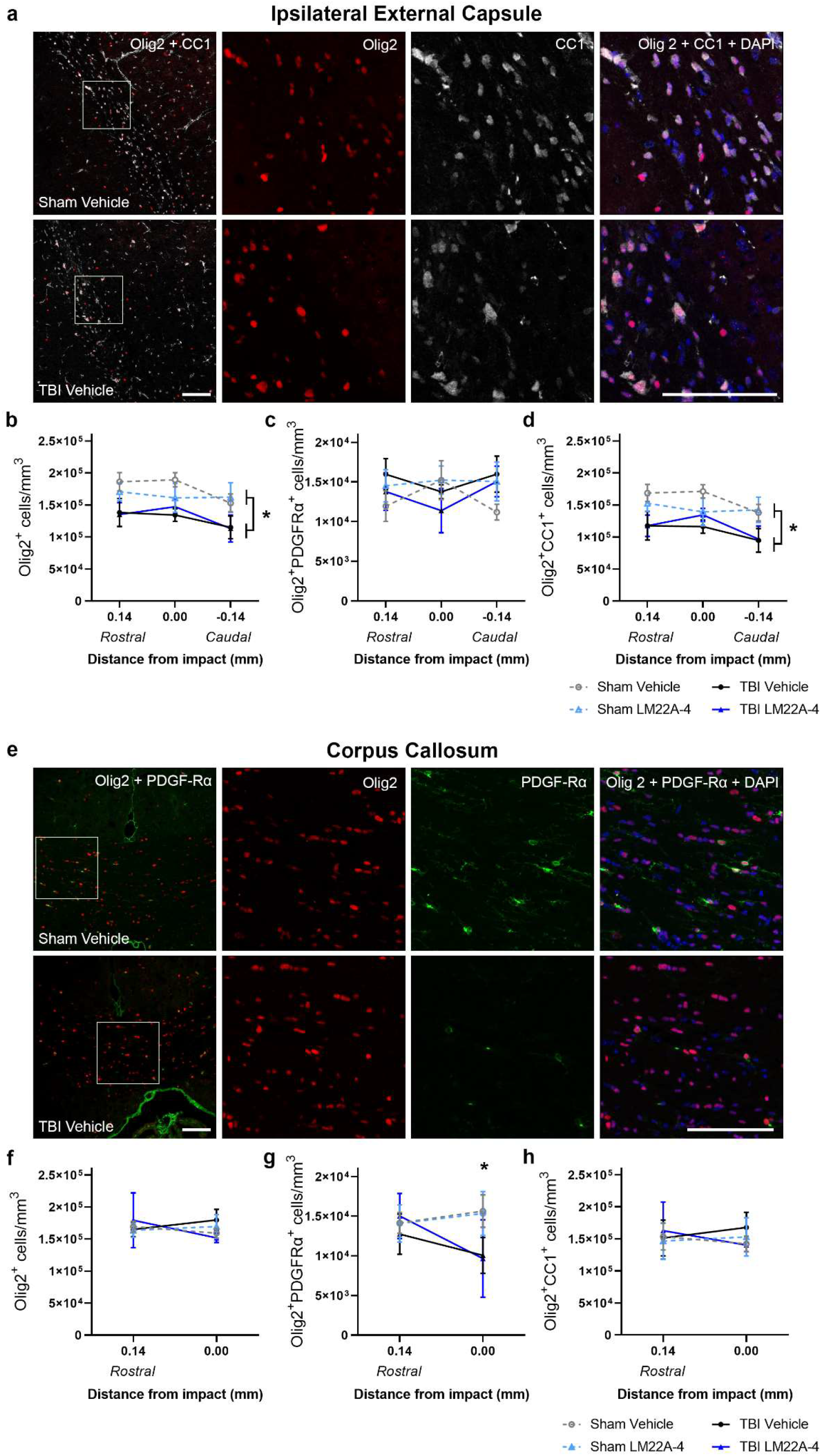
Loss of post-mitotic oligodendrocytes and oligodendrocyte progenitor cells (OPCs) persists up to 5 weeks post-TBI and is regionally dependent on distance from impact. Representative micrographs of Olig2 and CC1 co-immunolabeling (a), used to identify post-mitotic oligodendrocytes, in the ipsilateral external capsule of sham-vehicle and TBI-vehicle treated mice, showing reduced density of Olig2^+^ only and Olig2^+^CC1^+^ cells in TBI mice (*post-hoc p=0.013), as quantified in (b, d). LM22A-4 treatment had no effect. In the ipsilateral external capsule (c), the Olig2^+^PDGFRα^+^ OPC density was unchanged in sham and TBI mice (p=0.82), irrespective of treatment with LM22A-4 (p=0.98). Representative micrographs of Olig2 and PDGFRα co-immunolabeling (e), used to identify OPCs in the medial corpus callosum of sham-vehicle and TBI-vehicle treated mice, showing reduced density of Olig2^+^PDGFRα^+^ cells at the level of the impact site in TBI mice (*post-hoc p=0.007), as quantified in (g). The total density of Olig2^+^ cells (f), and Olig2^+^CC1^+^ cells (h) in the medial corpus callosum was unaltered by TBI or LM22A-4 treatment. For (a, e) scale bar = 100 μm. For (b-d, f-h), mean ± SEM plotted, n=6/group, REML, Bonferroni adjusted p-values used for post-hoc comparisons.

Oligodendroglial counts at 5 weeks post-TBI in the ipsilateral external capsule revealed that the density of Olig2^+^ cells followed a rostro-caudal gradient, with reduced density at 0.14 mm caudal from the impact site in all mice (REML Mixed model, distance F_1,2_= 11.14, p<0.001; Figure 3b). In TBI mice, irrespective of LM22A-4 treatment, the density of Olig2^+^ cells was reduced at the level of the impact site and in the adjacent rostral and caudal perilesional tissue when compared to sham-vehicle control mice (REML Mixed Model, injury, F_1,2_=6.99, p=0.016). The density of Olig2^+^PDGFRα^+^ OPCs was unchanged across the rostro-caudal axis of the ipsilateral external capsule (REML Mixed model, location, F_1,2_=0.44, p=0.64) in both sham-control and TBI mice, irrespective of injury (F_1,2_=0.051, p=0.82) or LM22A-4 treatment (F_1,2_=0.00049, p=0.98; Figure 3c). This is consistent with the characteristic uniform ‘tiled’ density of OPCs throughout the brain regardless of anatomical region (Hughes et al., 2013). Assessment of post-mitotic oligodendrocytes revealed a rostro-caudal decrease in the density of Olig2^+^CC1^+^ cells in sham-control and TBI mice at 0.14 mm caudal from the impact site (REML Mixed Model, distance, F_1,2_=11.63, p<0.001; Figure 3d). Importantly, compared to sham-vehicle control mice, there was an approximate 30-33% reduction in Olig2^+^CC1^+^ cell density at the level of the impact site, as well as rostral and caudal perilesional areas of TBI mice (injury, F_1,2_=7.47, p=0.013). LM22A-4 treatment had no effect on Olig2^+^CC1^+^ cell density in the ipsilateral external capsule of either sham or TBI mice (F_1,2_=0.73, p=0.79). Overall, these data from the ipsilateral external capsule are consistent with a long-term perturbation in either the survival, or potentially the differentiation, of post-mitotic oligodendrocytes in brain regions directly affected by TBI.

In the medial corpus callosum, the density of Olig2^+^ cells was unaffected by either the rostro-caudal axis (REML Mixed Model, distance, F_1,2_=1.48, p=0.23), TBI (F_1,2_=0.82, p=0.37), or LM22A-4 treatment (F_1,2_=0.41, p=0.53; Figure 3f). However, examination of Olig2^+^PDGFRα^+^ cells revealed that OPC density in the corpus callosum at the level of the impact site was reduced by approximately 45% following TBI compared to sham controls (REML Mixed Model, distance × injury interaction, F_1,2_=6.09, p=0.003), which was not ameliorated by LM22A-4 treatment (F_1,2_=0.62, p=0.44; Figure 3g). At 0.14 mm rostral to the impact site, the callosal OPC density was unaltered by TBI (Bonferroni’s post-hoc p=0.99; Figure 3g). Olig2^+^CC1^+^ post-mitotic oligodendrocytes demonstrated a trend towards an increased density in the corpus callosum of TBI mice at the level of the impact site (REML Mixed Model, distance × treatment × injury interaction, F_1,2_=2.79, p=0.07; Figure 3h). These data show that there are reduced OPCs in myelinated tracts adjacent to the site of impact that persists up to 5 weeks post-TBI, and is unchanged by acute treatment with LM22A-4. Overall, our findings from the corpus callosum and ipsilateral external capsule, indicate that—even 5 weeks after paediatric TBI—there persists regionally distinct disturbances to oligodendroglial sub-populations that are influenced by the distance from the impact site, but are unaffected by acute treatment with LM22A-4 immediately post-injury.

### Reactive gliosis persists at 5 weeks post-TBI, but is reduced by acute intranasal treatment with LM22A-4

TBI is accompanied by a robust neuroinflammatory response consisting of gliosis and chronic glial scarring (DiSabato et al., 2017), which was assessed by immunofluorescence staining in the bilateral external capsules and corpus callosum at 5 weeks post-TBI (Figure 4a). In the corpus callosum, evaluation of the percentage area positive for each marker revealed a trend towards increased GFAP^+^ astrogliosis (2way ANOVA, F_1,20_=3.30, p=0.08) and Iba1^+^ microgliosis (F_1,20_=3.77, p=0.07) in TBI mice, that were not altered by LM22A-4 treatment (data not shown).

**Figure 4:**
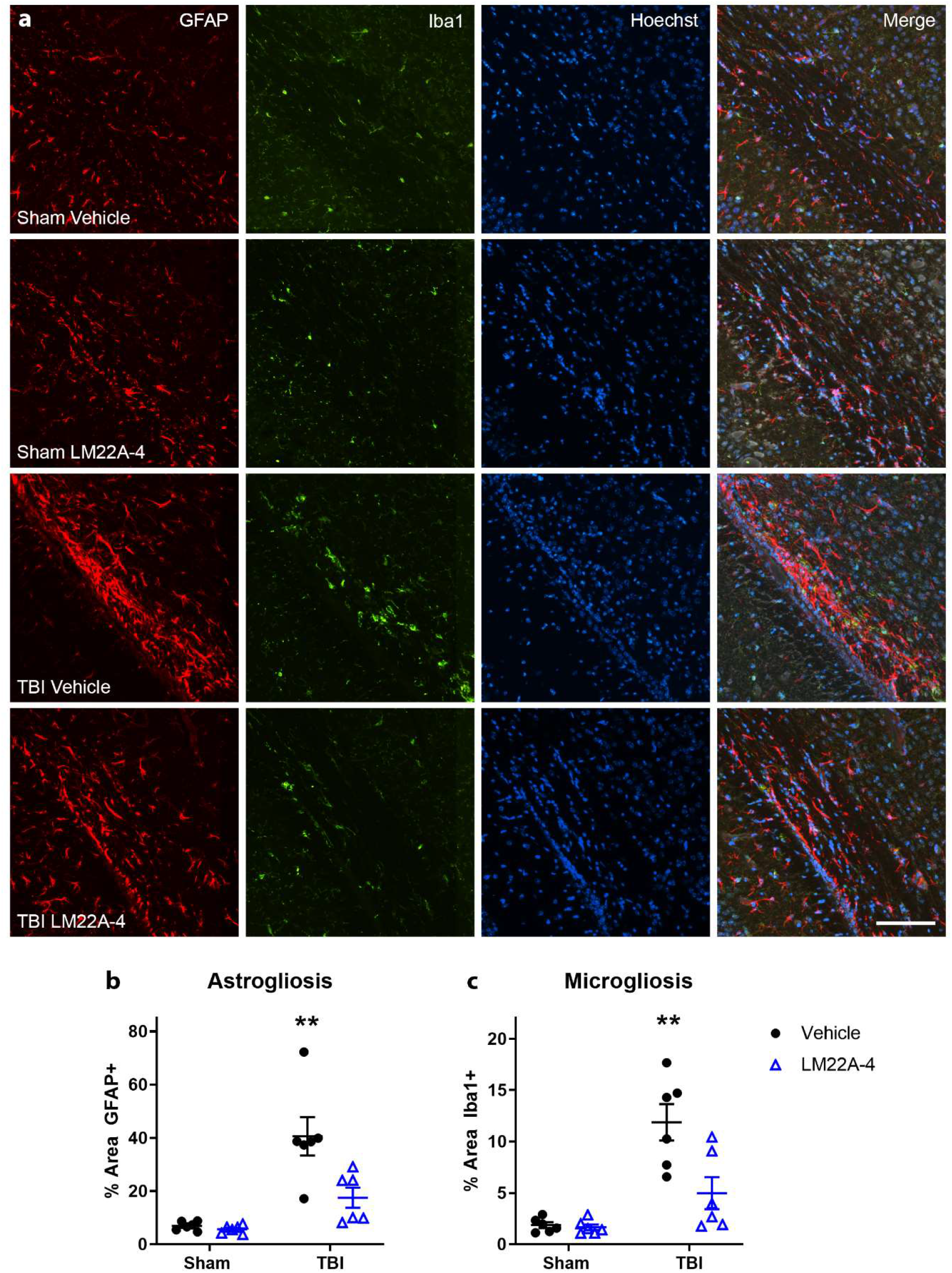
Glial reactivity in the ipsilateral external capsule at 5 weeks post-TBI is reduced by acute intranasal LM22A-4 treatment. Representative micrographs of GFAP and Iba1 co-immunolabeling (a) used to identify reactive astrocytes and microglia/macrophages respectively, in the ipsilateral external capsule of sham-vehicle, sham-LM22A-4, TBI-vehicle and TBI-LM22A-4 treated mice. Scale bar = 100 μm. The percentage area of positive GFAP (b) and Iba1 (c) staining in the ipsilateral external capsule was elevated in TBI-vehicle mice (both: *post-hoc p<0.001), but this increase was attenuated in TBI mice that received LM22A-4 treatment. For (b-c), mean ± SEM plotted, n=6/group, 2-way ANOVA, Tukey’s post-hoc test used for multiple comparisons.

Astrogliosis and microgliosis were pronounced in the ipsilateral external capsule of TBI mice, where increased GFAP^+^ and Iba1^+^ immunostaining was apparent compared to both sham-vehicle and LM22A-4-treated control mice (GFAP: 2-way ANOVA, injury × treatment interaction, F_1,20_=7.09, p=0.015, Tukey’s post-hoc p<0.0001, and Iba1: F_1,20_=7.80, p=0.011, post-hoc p<0.0001). Treatment with LM22A-4 commencing immediately post-TBI attenuated this gliosis, with a reduction in GFAP^+^ astrogliosis (Tukey’s post-hoc p<0.01; Figure 4b) and Iba1^+^ microgliosis (Tukey’s post-hoc p<0.01; Figure 4c).

### TBI results in cortical and myelinated tract atrophy, which is ameliorated by LM22A-4

Overt changes in brain morphology, including regional atrophy, are frequently observed after clinical TBI as well as in animal models, and may contribute to the manifestation of observed functional deficits (Osier et al., 2015). Therefore, to assess whether LM22A-4 influenced the extent of tissue damage after pediatric TBI, the volume of intact tissue in the dorsal cortex, hippocampus, corpus callosum and external capsule was quantified from Luxol fast blue or cresyl violet-stained sections (Figure 5a, d). In the dorsal cortex ipsilateral to the injury site, there was a reduction in cortical volume in TBI-vehicle animals specifically (2-way ANOVA, injury × treatment interaction, F_1,20_= 6.61, p=0.018), whereas TBI-LM22A-4 animals had a similar cortical volume to sham control mice (Bonferroni’s post-hoc between TBI groups, p<0.01; Figure 5b). In the ipsilateral hippocampus, neither TBI nor LM22A-4 treatment influenced intact tissue volume (2-way ANOVA, treatment, F_1,20_=1.78, p=0.20; injury, F_1,20_=0.88, p=0.86; Figure 5c).

**Figure 5:**
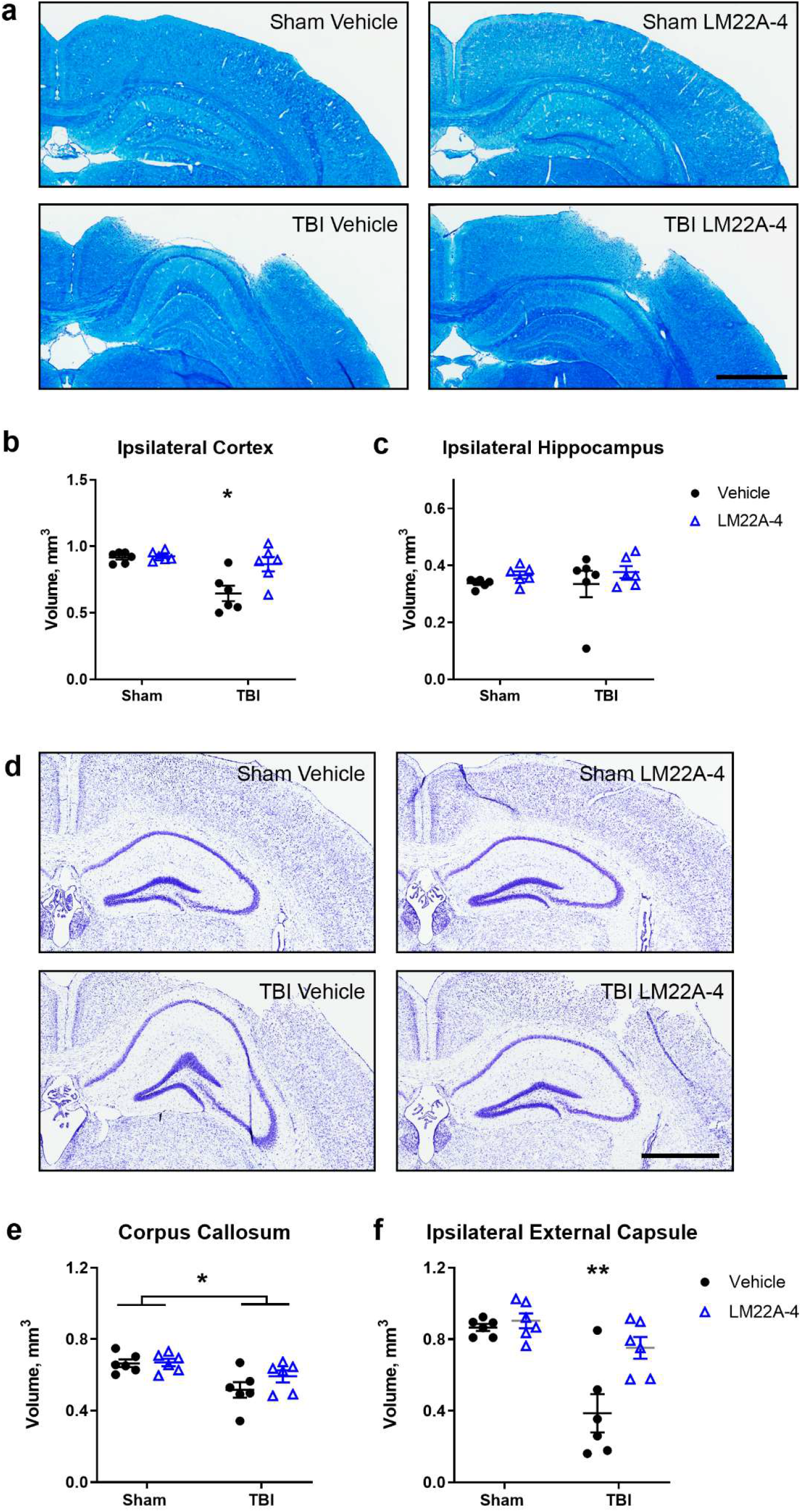
TBI-induced atrophy of the cortex and ipsilateral external capsule is prevented by acute intranasal treatment with LM22A-4. Representative micrographs of Luxol Fast Blue staining of coronal brain sections (a) were used to delineate cortical and hippocampal tissue volumes at 5 weeks post-injury. The ipsilateral cortical volume (b) was decreased at 5 weeks post-injury in TBI-vehicle mice compared to those that received LM22A-4 (*post-hoc p<0.01), but there was no injury effect on hippocampal volumes (c). Representative micrographs of cresyl violet stained coronal brain sections (d) used to quantify tissue volume in the corpus callosum and ipsilateral external capsule. In the corpus callosum (e), tissue volume was decreased with TBI (*p=0.0018) and not preserved by LM22A-4 treatment. However, the TBI-induced tissue volume loss in the ipsilateral external capsule (f) was prevented in TBI mice that received LM22A-4 (*post-hoc p=0.0042). For (a, d) scale bar = 1000 μm. For (b-c, e-f), mean ± SEM plotted, n=6/group, 2-way ANOVA, with Tukey’s post-hoc test used for multiple comparisons.

The volume of the corpus callosum was reduced at 5 weeks post-TBI compared to both sham-control groups (2-way ANOVA, F_1,20_=12.95, p=0.0018). However, in contrast to the dorsal ipsilateral cortex, LM22A-4 treatment after TBI did not preserve callosal volume (F_1,20_=1.65, p=0.21; Figure 5e). In the ipsilateral external capsule, intact tissue volume was reduced in TBI-vehicle mice compared to sham control mice (2-way ANOVA treatment × injury interaction, F_1,20_=6.24, p=0.021; Tukey’s post-hoc p=0.0003). Of note, external capsule volume was preserved in TBI mice that received LM22A-4 compared to TBI-vehicle mice (p=0.0042), with TBI-LM22A-4 treated mice having similar volumes to sham controls (p=0.386; Figure 5f). The volumes of neuroanatomical regions contralateral to the injury site (dorsal cortex, hippocampus and external capsule) were not affected by either injury or LM22A-4 treatment (data not shown). Collectively, these data indicate that intranasal delivery of LM22A-4 immediately post-injury improves brain atrophy outcomes by reducing tissue volume loss in brain regions proximal to the injury site.

### Treatment with LM22A-4 improves TBI-induced hyperactivity and anxiolytic behaviors, but does not influence motor function or social memory impairments

Between 4-5 weeks post-injury, mice underwent a battery of neurobehavioral tests to evaluate sensorimotor, cognitive and psychosocial function. In the open field arena (Figure 6a), as expected based on previous findings of hyperactivity in this model of pediatric TBI (Semple et al., 2014b), brain-injured mice were observed to travel a greater distance in this task (2-way ANOVA effect of injury F_1,20_=2.51, p=0.0002). Interestingly, LM22A-4 treatment reduced the distance moved by both the sham-injured and TBI mice (effect of treatment, F_1,20_=5.35, p=0.03).

**Figure 6:**
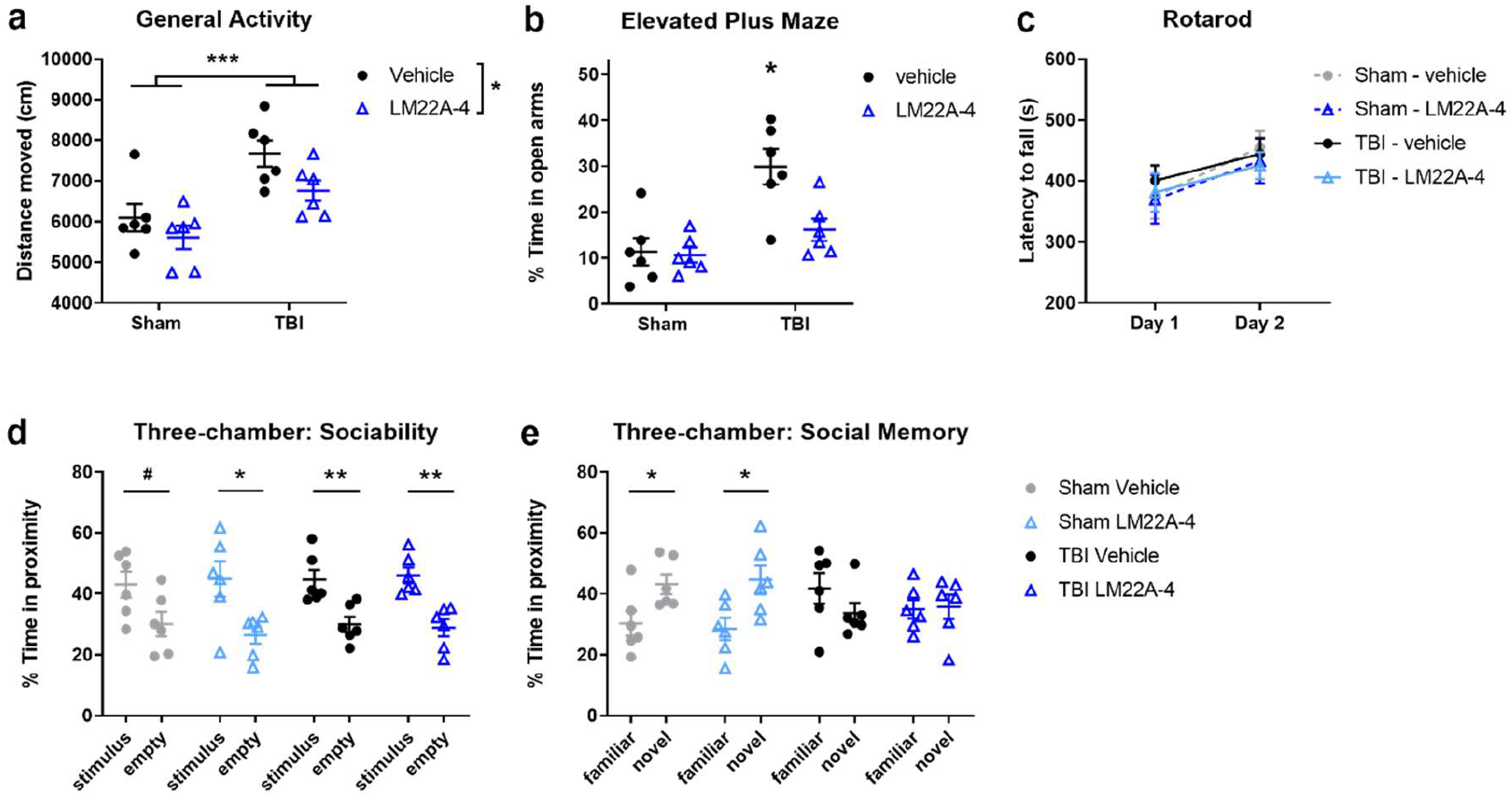
LM22A-4 treatment improves TBI-induced hyperactivity and anxiolytic behaviors but does not improve social memory impairments that emerge 4-5 weeks postinjury. In the open field (a), TBI overall increased general activity (***p=0.0002) while treatment with LM22A-4 reduced this (*p=0.03). In the Elevated Plus Maze (b), TBI-vehicle mice spent more time in the open arms (*p=0.03) compared to both groups of sham animals and the LM22A-4 treated TBI mice. All groups improved over time in the rotarod task (c). In the three-chamber test for sociability (d), all groups showed a preference for the chamber with the stimulus mouse, compared to an empty chamber (#p=0.052; *p<0.05; **p<0.01). In the subsequent stage of the task (e), to evaluate social memory, mice in the sham groups showed a preference for a novel stimulus mouse over a now-familiar stimulus mouse (*p<0.05), while TBI injured mice failed to show this preference even with LM22A-4 treatment. For (a-e), mean ± SEM plotted, n=6/group. For (a, b) 2-way ANOVA, (c) 3-way repeated measures ANOVA, both with Tukey’s post-hoc test for multiple comparisons. For (d, e) unpaired t-test.

In the Elevated Plus Maze, TBI-vehicle mice spent an average of 30% of their time in the open arms, compared to approximately 10% in sham control mice (Figure 6b), indicative of reduced anxiety-like behaviour after TBI consistent with previous observations in this pediatric TBI model (Pullela et al., 2006). Indeed, time spent in the open arms by TBI-vehicle mice was also increased compared to LM22A-4 treated TBI mice (2-way ANOVA, injury × treatment interaction, F_1,20_=5.26, p=0.03), which spent approximately 10-15% of their time in the open arms. These findings suggest that treatment with LM22A-4 immediately after TBI alleviates the anxiolytic effects of injury at this time point.

The rotarod was performed as a measure of gross motor function, coordination, and motor learning, across two consecutive days (Figure 6c). While performance was improved in the task between days 1 and 2 (3-way repeated measures ANOVA, time, F_1,20_=7.63, p=0.01), neither TBI (F_1,20_=0.04, p=0.84) nor LM22A-4 treatment (F_1,20_=0.52, p=0.48) resulted in any change to motor function and learning at this time.

Finally, the three-chamber social approach test has previously been demonstrated to be sensitive to deficits in social interest and social recognition memory, such as those exhibited chronically more than 8 weeks after pediatric TBI (Semple et al., 2012; Semple et al., 2017). Here, following habituation to the apparatus, mice were first presented with a choice between outer chambers containing either a novel stimulus mouse or an empty cage to examine their preference for sociability (Figure 6d). Most groups exhibited a preference for the stimulus-containing chamber compared to the empty chamber (unpaired t-tests: sham-LM22A-4 t-test p=0.017; TBI-vehicle p=0.006; TBI-LM22A-4 p=0.001), while sham-vehicle mice also demonstrated a trend towards this preference (unpaired t-test: p=0.052). This indicates that behavior was typically intact in TBI mice up to 4-5 weeks post-injury, and unaffected by LM22A-4 treatment.

In the subsequent test stage, a second novel stimulus animal was placed within the previously empty chamber, providing the experimental animal with a choice between a now-familiar stimulus and a novel stimulus. Social recognition, or social memory, is indicated by a preference for the novel stimulus mouse over the now-familiar stimulus mouse (Figure 6e). In this task, both sham groups showed a preference for the chamber containing the novel mouse (unpaired t-tests: sham-vehicle p=0.034; sham-LM22A-4 p=0.022). In contrast, neither TBI group showed such a preference (TBI-vehicle p=0.207; TBI-LM22A-4 p=0.882). These findings indicate that pediatric TBI results in social memory impairments that can be detected 4-5 weeks post-injury, and are not alleviated by acute LM22A-4 treatment.

Overall, these behavior data are suggestive that neuroprotection and preservation of myelin integrity mediated by acute treatment with the partial TrkB agonist LM22A-4 can improve TBI-associated hyperactivity and anxiolytic behaviors, but there is no influence on social memory impairments.

## DISCUSSION

Monitoring of lesion progression after TBI by MRI has resulted in focused attention on how damage to myelinated tracts after early life brain injuries contribute to neurobehavioral dysfunction in adulthood (Gale and Prigatano, 2010; Genc et al., 2017; Zamani et al., 2020). Importantly, the cellular and molecular mechanisms underlying these MRI measures in the context of pediatric TBI—where ongoing neurodegenerative damage is superimposed on active postnatal and adolescent neurodevelopment—are poorly understood. Using a mouse model of pediatric TBI, here we identified acute damage to compact myelin in the injured hemisphere, resulting in persistent fragmentation and structural perturbations of compact myelin that continues for at least 5 weeks post-injury. Loss of compact myelin was limited to directly affected tissue regions and accompanied by TBI-induced deficits in oligodendroglial populations, reactive gliosis, atrophy of the corpus callosum and external capsule, and neurobehavioral changes.

Intriguingly, only dysmyelination of the perilesional external capsule was normalized to healthy levels by intranasal treatment with the partial TrkB agonist, LM22A-4. This was accompanied by reduced atrophy in the ipsilateral dorsal cortex and external capsule, and reduced localized gliosis, suggesting that the neuroprotective properties of LM22A-4 (Massa et al., 2010) are more important in the context of pediatric TBI than its reported pro-myelinating activity (Geraghty et al., 2019; Nguyen et al., 2019). LM22A-4 treatment also ameliorated TBI-associated hyperactivity and anxiolytic behaviors that emerged during adolescence, but had no effect on social memory impairments. Notably, impaired sociability and social recognition have previously been reported to develop with age, and manifest in adult pediatric brain-injured mice aged at least 2 weeks older than the mice examined here (Semple et al., 2012; Semple et al., 2014a). This time-dependent emergence of neurobehavioral dysfunction emphasizes the need for longitudinal intervention studies which examine efficacy of novel treatments on both cellular pathophysiology and chronic behavioral outcomes in pediatric TBI.

Our study demonstrates that acute structural damage to myelinated tracts during rapid and ongoing developmental myelination results in persistent myelin fragmentation and loss, that continues up to 5 weeks post-injury (i.e. 8 weeks of age), when brain-injured mice are reaching early adulthood (Semple et al., 2013). These changes are most pronounced at the site of injury, in the proximal external capsule, likely as a consequence of both primary mechanical damage as well as secondary neurodegeneration. However, reduced levels of myelination were found to extend across the entire perilesional zone, indicating that this pathology likely has widespread and profound impacts on neural circuit function, as demonstrated by the hyperactive and anxiolytic behavior exhibited by vehicle-treated TBI mice. Axonal damage and loss due to TBI are the likely primary drivers of these myelin changes. However, the lack of detectable βAPP staining at 5 weeks post-injury renders it difficult to determine the relative contribution of underlying axonal injury to myelin loss and dysmyelination in this model versus the influence of intrinsic oligodendroglial dysfunction, or an inhibitory inflammatory microenvironment resulting from the TBI. This limitation could be overcome in future studies by the use of a Thy1 fluorescent reporter strain to endemically label axons, an approach that has been used successfully by others to assess focal and diffuse axonal injury (Marion et al., 2018; Nikić et al., 2011).

Both post-mitotic oligodendrocytes and OPCs were lost following pediatric TBI, as anticipated, although the regional and sub-population heterogeneity of this cellular response was unexpected based on previous studies in mild and severe TBI to the adult brain (Dent et al., 2015; Mierzwa et al., 2015; Mierzwa et al., 2014). However, there is currently no true consensus on how oligodendroglia respond to TBI in either the adult or pediatric brain, other than an acute increase in OPC proliferation (Dent et al., 2015; Flygt et al., 2013; Mierzwa et al., 2015; Warnock et al., 2020). We did not assess cell proliferation in the current study, due to our unique focus on the pediatric context and medium-term recovery phase. Instead, a reduction in OPC density was observed at 5 weeks post-injury in the corpus callosum, distal but adjacent to the injury site, and no OPC changes in the injured external capsule. Our use of triple-labeling indicates that the reduction in OPC density is not due to an overall loss of oligodendroglia—indeed, based on corresponding increases in Olig2^+^CC1^+^ cells in TBI animals, this finding is more likely attributed to a subtle acceleration in differentiation. OPC migration, to replace oligodendrocytes in injured regions, cannot be definitively excluded as a possibility; however, this seems unlikely as OPCs have a limited migration range (Hughes et al., 2013).

At 5 weeks following pediatric TBI, a reduced density of Olig2^+^CC1^+^ oligodendrocytes was observed in the ipsilateral external capsule. This is in contrast with previous studies in adult models of TBI, where there were no prolonged effects on the density of post-mitotic oligodendrocytes (Dent et al., 2015; Mierzwa et al., 2015; Mierzwa et al., 2014). In a comparable controlled cortical impact model of experimental TBI, in adult injured mice, an acute loss of CC1^+^ oligodendroglia was reported in the injured external capsule, that returned to sham-control levels by 5 weeks post-injury, while there were no oligodendroglial density changes in the corpus callosum (Dent et al., 2015). Intriguingly, this previous study also reported an elevation of CC1^+^ cells co-labeled with the apoptosis marker activated caspase-3 at 5 weeks post-TBI, indicative of ongoing intrinsic dysfunction causing the progressive loss of these cells in the injured external capsule. This suggests that intracellular Ca^2+^ overload, oxidative stress, and lipid peroxidation—all mechanisms proposed to drive oligodendroglial death in neurotrauma—propagate chronically after TBI to the adult brain (Warnock et al., 2020). At face-value, our results of reduced post-mitotic oligodendrocytes 5 weeks post-injury, indicate that these pathomechanisms are also chronically active in pediatric TBI, contributing to decreased oligodendrocyte survival. However, as our single time point did not allow for evaluation of whether or when cell numbers return to sham-control levels, there could be an additional impairment in the differentiation of post-mitotic oligodendrocytes after TBI that is specific to the pediatric context, and may relate to the effects of axonal dysfunction and loss during late postnatal brain development. Both this, and the intriguing reduction in OPC density in the corpus callosum, could be resolved by future fate-mapping studies.

Our key finding in this study was that acute intranasal treatment with the partial TrkB agonist LM22A-4 preserved (or repaired) myelin integrity, reduced reactive gliosis, prevented cortical and myelinated tract atrophy, and ameliorated hyperactive and anxiolytic behaviors in TBI mice, but it did not preserve oligodendrocyte populations. LM22A-4 was used in this study with the specific hypothesis that it would target myelin repair, as oligodendroglial BDNF-TrkB signaling has previously been shown to promote myelination via the MAPK/Erk pathway in oligodendrocytes (Du et al., 2006; Van’t Veer et al., 2008). Further, LM22A-4 has been successfully used to promote oligodendrogenesis, as well as increase oligodendrocyte differentiation and the extent of myelin repair in an adult mouse model of cuprizone toxin-induced demyelination (Nguyen et al., 2019). From this perspective, the dissonance between reduced Olig2^+^CC1^+^ cells and preserved myelin integrity in the injured external capsule of LM22A-4 treated TBI mice is difficult to reconcile, despite the obvious contextual differences of disease model, LM22A-4 dose, delivery mode and treatment length, as we have recently observed an 1.6-fold increase in Olig2^+^CC1^+^ cells mediated by LM22A-4 treatment (via oligodendroglial TrkB) in the cuprizone study (Nguyen et al., 2019). This finding was further supported by independent studies utilizing LM22A-4 in the lysolecithin model of demyelination (Geraghty et al., 2019). Further investigation is required to determine whether LM22A-4 in our hands indeed resulted in an acute increase in oligodendroglia (e.g. during or after the 14 days of treatment) which may have not survived until the 5 week post-injury time point evaluated here.

Regardless, BDNF-TrkB signaling has been identified as a potential therapeutic target for over 25 years (Spina et al., 1992), primarily driven by its early and well characterized roles in neuronal survival and synaptic plasticity (Chao, 2003). Importantly, LM22A-4 has been used both intranasally and intraperitoneally in the context of adult TBI, where it resulted in improved motor learning 2-3 weeks after injury (Massa et al., 2010). However, neither behavioral nor histological assays were undertaken in this previous study, such that the mechanisms of action were not defined. More recently, LM22A-4 has been used in a partial neocortical isolation model of trauma-induced epilepsy in adult rodents (Gu et al., 2018). In this context, LM22A-4 prevented cortical hyperexcitability, suppressing post-traumatic epileptogenesis and increased putative inhibitory synapses (Gu et al., 2018). Together, our current findings provide further insight into the mechanism of LM22A-4 action in neuroprotection, by both preserving myelin integrity and cortical and external capsule tissue volumes, as well as ameliorating hyperactive and anxiolytic behaviors after pediatric TBI. Hyperactivity and anxiolytic behaviors induced by TBI are associated with cortical volume loss and loss of dendrite complexity (Pullela et al., 2006; Semple et al., 2017). BDNF-TrkB signaling is well established to influence dendrite morphology and stabilize synapses (Gorski et al., 2003; Lin and Koleske, 2010; Tolwani et al., 2002), and it is possible that LM22A-4 via TrkB is mimicking this effect here to preserve tissue volume and neural circuit function.

Critically, if LM22A-4 does improve circuit function post-TBI, it also raises the possibility that activity-dependent myelination mechanisms (Suminaite et al., 2019) are required to preserve myelin post-injury, and that the scenario is more complicated than simply preserving the axonal substrate for oligodendrocytes to myelinate, as currently assumed (Mierzwa et al., 2015). This line of thought also raises critical questions about the pathogenesis of social memory impairments in male pediatric TBI mice, which here was demonstrable at 4-5 weeks postinjury, but was not responsive to LM22A-4 treatment. Previously, we hypothesized that corpus callosum atrophy is involved in social behavior deficits after pediatric TBI (Semple et al., 2012; Semple et al., 2014a). Aligned with this view of subtle changes in structural connectivity triggering social impairments after TBI—even though the level of myelination was not reduced compared to sham controls—the natural rostro-caudal variation in myelination of the corpus callosum was *not* observed in TBI mice, regardless of treatment. Of course, this speculation requires formal examination via longitudinal studies that monitor the progression of social impairments, combined with *in vivo* imaging modalities such as DTI and *ex vivo* histological assessment (Zamani et al., 2020). Overall, our results speak to the multi-modal neuroprotective effects of TrkB signaling in the CNS, and indicate that in this specific context of pediatric TBI, the improved myelin outcomes are likely due to a TrkB-dependent effect on preserving neuronal function. This is in contrast to a hypothesized direct effect on oligodendroglial TrkB, which could be further defined by using neuronal and oligodendroglial TrkB conditional knockout mice.

## CONCLUSION

Our study demonstrates that treatment with a partial TrkB agonist immediately following TBI in pediatric mice preserves myelin integrity, reduces reactive gliosis and prevents cortical and myelinated tract atrophy in the medium-term recovery phase. These improved neurobiological outcomes due to acute treatment with LM22A-4 likely contributed to the amelioration of hyperactive and anxiolytic behaviors in treated mice 5 weeks after pediatric brain injury. However, persistent oligodendroglial density reductions and social memory defects did not respond to LM22A-4 treatment. Overall, our findings suggest that the multi-modal neuroprotective effects of TrkB signaling drove the positive outcomes of acute LM22A-4 treatment, and provide promise that acute pharmacological intervention can modulate both pathological and neurobehavioral outcomes after pediatric TBI.

## CRediT author statement

Jessica Fletcher: Conceptualization, methodology, software, validation, formal analysis, investigation, resources, data curation, writing – original draft, review and editing, supervision, project administration, funding acquisition. Larissa Dill: Methodology, software, validation, formal analysis, investigation, data curation, writing – review and editing, visualization. Rhiannon Wood: Investigation, validation. Sharon Wang: Investigation, formal analysis. Kate Robertson: Investigation, formal analysis. Simon Murray: Conceptualization, methodology, supervision. Akram Zamani: Methodology, validation, formal analysis, investigation, supervision. Bridgette Semple: Conceptualization, methodology, validation, formal analysis, investigation, resources, writing – original draft, review and editing, visualisation, supervision, project administration, funding acquisition.

## Acknowledgements

This work was supported by the National Health and Medical Research Council of Australia (NHMRC) via a Career Development Fellowship and Project Grant to BDS, as well as project funding from the Department of Anatomy and Neuroscience, The University of Melbourne, to JF and BDS. JF was supported by a 2018 Melbourne Neuroscience Institute Fellowship. Graphical abstract produced with BioRender©. The authors also thank Dr. David Gonsalvez (School of Biomedical Sciences, Monash University) for assistance with SCoRe imaging, as well as the Monash Histology Platform and Monash Micro Imaging Platform at the Alfred Research Alliance and the Biological Optical Microscopy Platform (BOMP) at the University of Melbourne for their facilities and technical support.

