## Supplementary Materials for "Acute treatment with TrkB agonist LM22A-4 confers neuroprotection and preserves myelin integrity in a mouse model of pediatric traumatic brain injury"

**Supplementary Methods: Fresh tissue collection**

For western blots, mice were transcardially perfused with 1x isotonic PBS before decapitation to remove the brain. The cerebral hemispheres and bilateral olfactory bulbs were dissected and flash-frozen in liquid nitrogen, then homogenized in ice cold lysis buffer (10 mM Tris, 150 mM NaCl, 1% lgepal in distilled water) containing phosphatase inhibitors (PhosSTOP Roche, 50 mM NaF) followed by centrifugation at 12,000 rpm. Total protein concentrations were determined by Bradford Assay (Sigma Aldrich). For Western blotting, 50 μg of protein was combined with Laemmli sample buffer for denaturation at 95 °C, then loaded into 12 well pre-cast protein gels (Bolt 4-12% Bis-Tris Plus, Invitrogen). Samples underwent sodium dodecyl sulfate-polyacrylamide gel electrophoresis at 200 V and 100 mA (~ 2 h), then transferred onto a polyvinylidene difluoride membrane (Trans-Blot Turbo Transfer Pack, Bio-Rad), using a Transblot Turbo Transfer System (Bio-Rad).

The membrane was washed in 1X Tris-buffered saline containing 0.5% Polysorbate-20, blocked with 2% bovine serum albumin to block non-specific binding, then incubated overnight at 4 °C with various primary antibodies against total TrkB (anti-rabbit-TrkB H-181, 1:1000; Santa Cruz), phosphorylated TrkB (anti-rabbit-pTrkB S478/479, 1:1000; Biosensis), total ERK1/2 (anti-rabbit-ERK1/2, 1:1000; Cell Signaling Technology) and phosphorylated ERK1/2 (anti-rabbit-pERK1/2, 1:1000; Cell Signaling Technology). The following day, membranes were washed then probed with an anti-rabbit IgG HRP-conjugated secondary antibody (1:5000; Cell Signaling Technology) for 2 h, then washed again. Protein bands were visualized on a ChemiDoc XRS + Imaging System using Clarity Western ECL Blotting Substrates (Bio-Rad) and automated exposure times, and bands were quantified as optical density from 8-bit images using FIJI Image J (Version 2.0.0). Presented blots are representative of at least 3 independent experiments.

**Supplementary Table 1: Experimental groups.**

| **Experiment** | **Time point** | **Aim** | **Sham** | **TBI** |
| --- | --- | --- | --- | --- |
| 1 | 3 days | Assess acute post-injury neuropathology | 3 | 3 |
| 2 | 2 weeks | Determine whether LM22A-4 induced TrkB signaling | 3 (vehicle)  3 (LM22A-4) | 3 (vehicle)  3 (LM22A-4) |
| 3 | 5 weeks | Evaluate chronic behavioral and pathological effects of LM22A-4 | 6 (vehicle)  6 (LM22A-4) | 6 (vehicle)  6 (LM22A-4) |

**Supplementary Table 2: Grading of MBP immunofluorescence staining.**

| **MBP Grade** | **Observations** |
| --- | --- |
| 0 | No evidence of myelin fragmentation  Normal-appearing structural integrity of myelinated tract |
| 1 | Minimal myelin fragmentation (< 3 distinct, intense MBP-bright particles)  Normal structure of myelinated tract |
| 2 | Considerable myelin fragmentation (>3 MBP-bright particles)  Normal structure of myelinated tract |
| 3 | Abundant myelin fragmentation and abnormal structure – narrowed, clearly-disrupted tract |

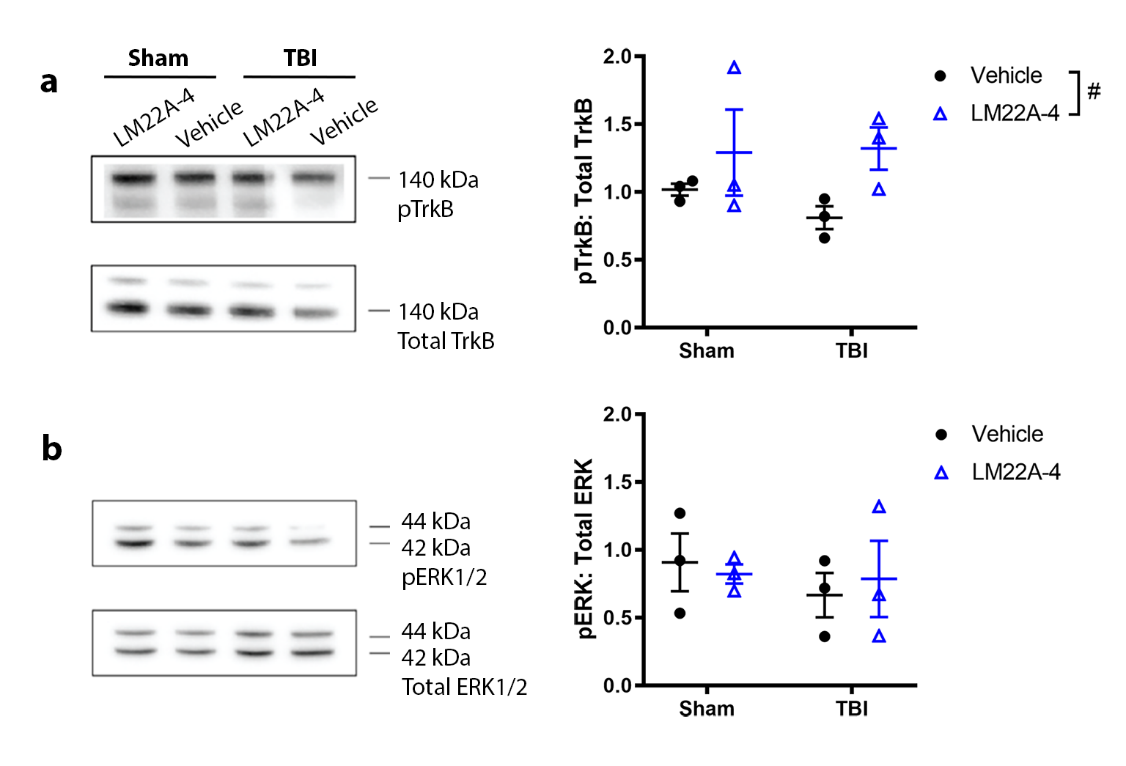

**Supplementary Figure 1: LM22A-4 effects on TrkB and ERK1/2 levels in the brain.** Using Western blotting, the presence of pTrkB, TrkB, pERK1/2 and ERK1/2 were detected in olfactory bulb and cerebral hemisphere homogenates at 14 days post-TBI or sham (n=3/group). In the olfactory bulbs (a), mice treated with LM22A-4 displayed a trend towards increased pTrkB (2-way ANOVA effect of treatment, F_1,8_=4.57, #p=0.06), independent of injury (F_1,8_=0.23, p=0.64). No differences in the level of phosphorylated ERK1/2 were observed in the olfactory bulbs (b). In the cerebral hemispheres, neither pTrkB or pERK1/2 were altered by injury or LM22A-4 treatment at this time point (not shown).

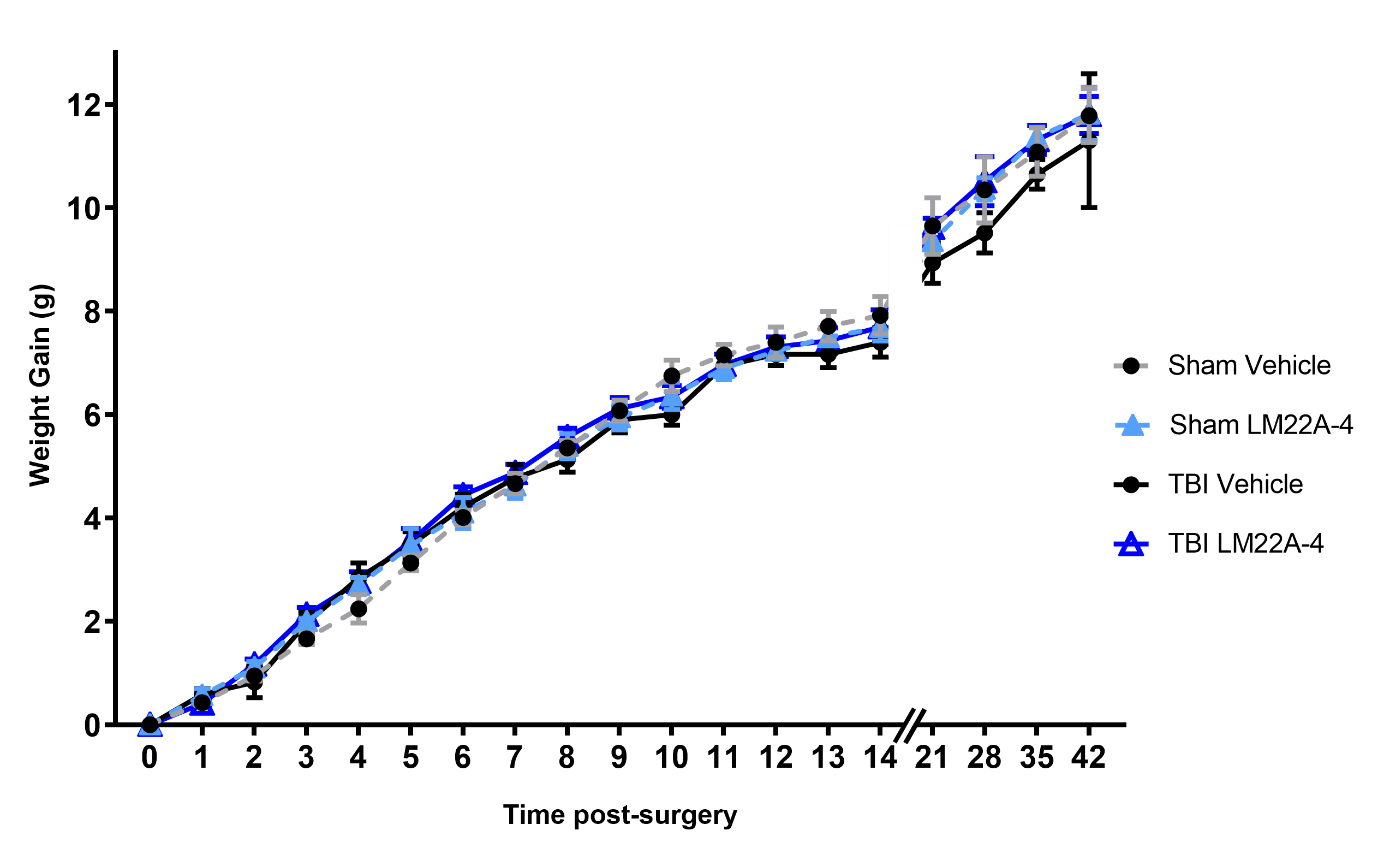

**Supplementary Figure 2: Body weights were not affected by LM22A-4 treatment.** No differences were observed between the groups at the time of surgery. Over time post-TBI or sham, all groups demonstrated an increase in body weight as anticipated (2-way ANOVA effect of time, F_218,360_=574.3, p<0.0001), but no difference between groups based on either TBI surgery or LM22A-4 treatment (F_3, 20_=0.51, p=0.68).

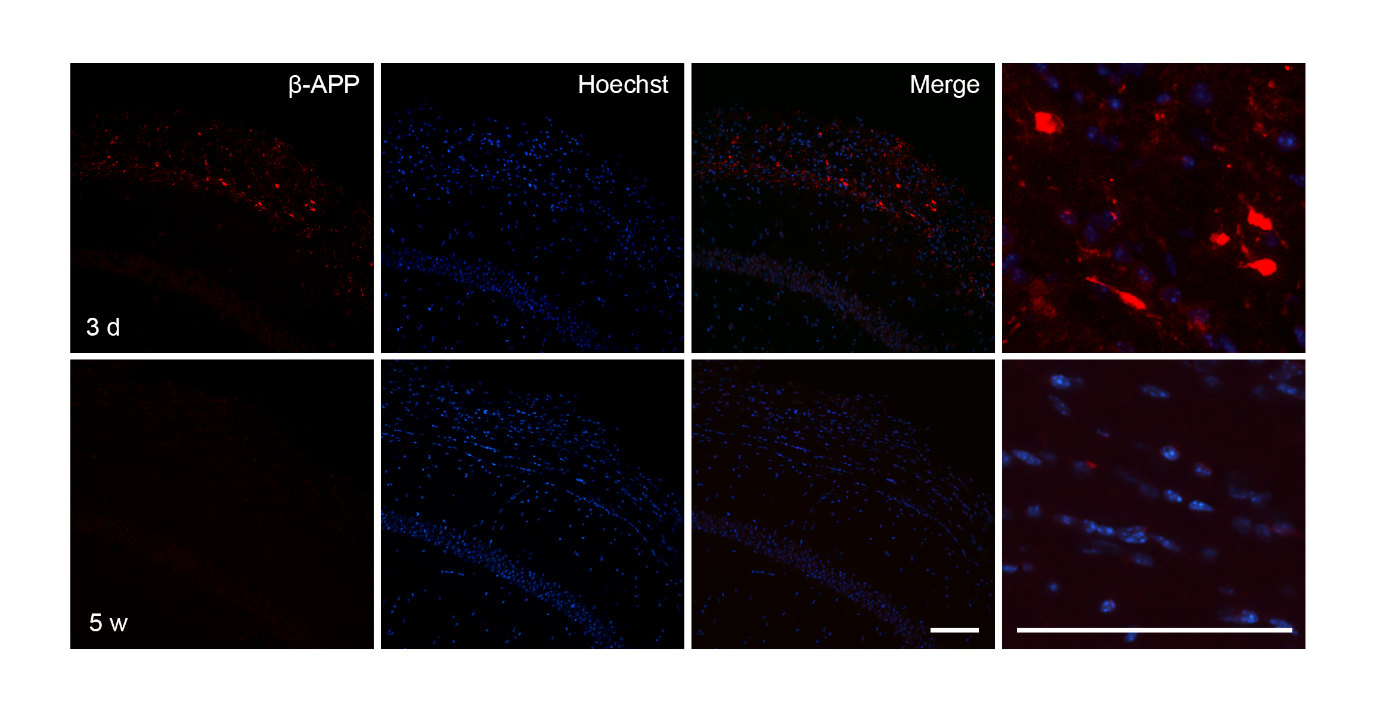

**Supplementary Figure 3: βAPP was minimally detected at 5 weeks post-TBI.** βAPP, an indicator of axonal injury, was immunolabeled after pediatric TBI. However, minimal positive staining was detected in the injured cortex, hippocampus or external capsule at 5 weeks post-TBI (lower panels). A 3 day time point (repeat of Figure 1) is also presented for comparison, and was used as a positive control specimen. Here, βAPP^+^ staining can be observed in the external capsule and injured cortex. Scale bars = 100 µm.

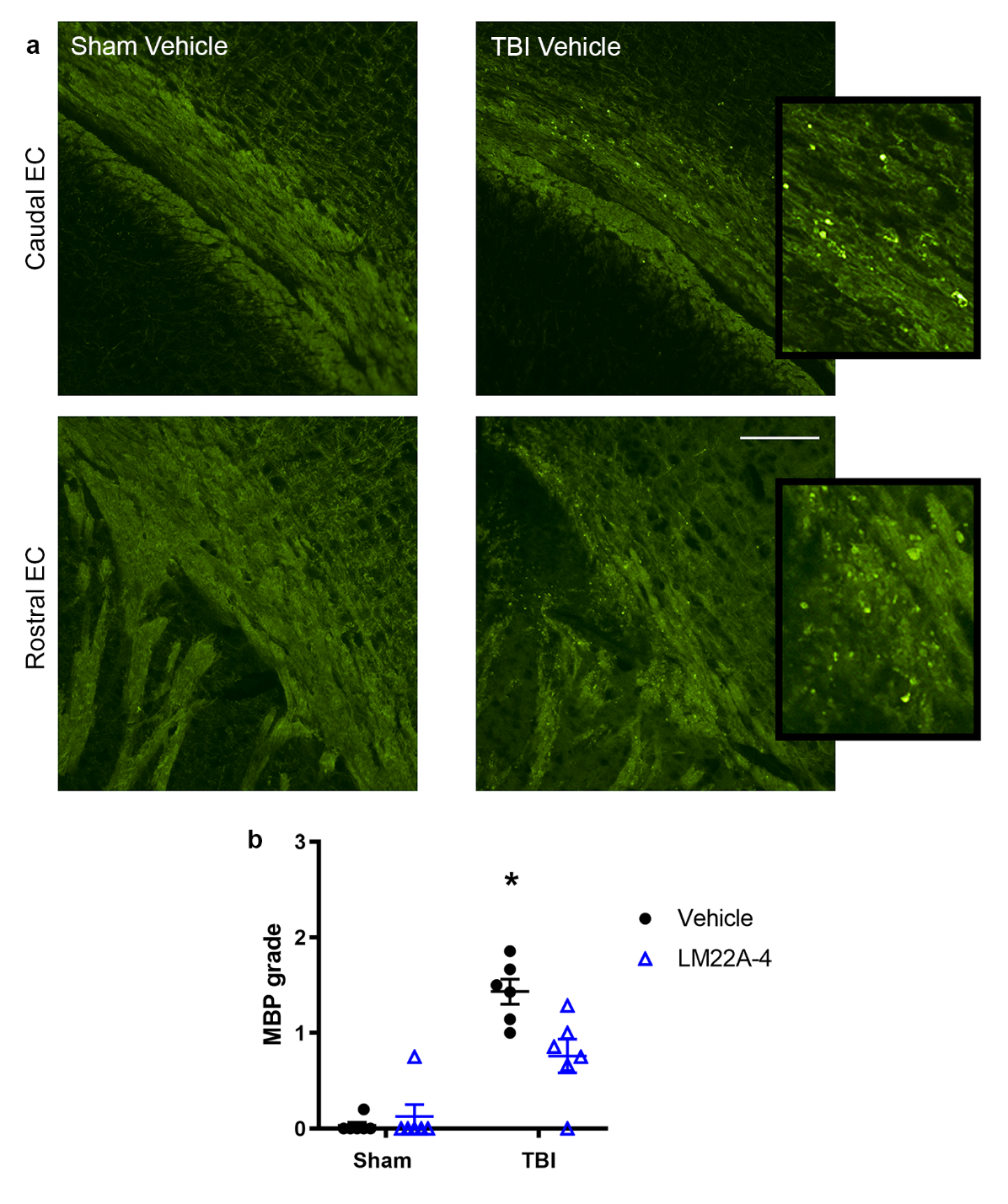

**Supplementary Figure 4: Pathology in myelinated tracts at 5 weeks post-TBI, detected by MBP immunofluorescence staining.** MBP immunofluorescence (a) revealed disruption of myelinated tracts in the ipsilateral external capsule of TBI mice. Qualitative assessment of this staining (b) confirmed evidence of pathology in the external capsule in TBI-vehicle mice, which was reduced in TBI-LM22A-4 mice (2-way ANOVA treatment $\times$ injury interaction F_1,20_=9.02, p=0.007; Bonferroni’s post-hoc, *p$\leq$0.05 vs. all other groups). Grading was performed by an investigator blinded to treatment/injury group as per the criteria in Supplementary Table 2, and the average grade of 4 sections per brain were compared. Mean ± sem plotted.
